# The Moment Symmetry Breaks: Spatiotemporal Dynamics of CYCLOIDEA Expression During Early Floral Development

**DOI:** 10.1101/2025.08.05.668671

**Authors:** Ya Min, Bianca T. Ferreira, Yao-Wu Yuan

## Abstract

The establishment of bilateral symmetry in flowers depends on the precise regulation of *CYCLOIDEA* (*CYC*) gene expression along the dorsal-ventral axis. Although auxin and *BLADE-ON-PETIOLE* (*BOP*) have been implicated as regulators of *CYC*, when, where, and how they affect *CYC* expression remains unclear. Here, we combined transgenic manipulation and fluorescent confocal imaging to capture the spatiotemporal dynamics of the *Mimulus parishii CYC* genes (*MpCYC2a* and *MpCYC2b*) in relation to FM growth and auxin activity maxima in both wild type (WT) and *bop* mutants (*mpbop*). Strikingly, *MpCYCs* have already gained dorsal expression in the FM prior to any detectable auxin maxima, and no difference in *MpCYC* expression was observed between the wild type and *mpbop* during this initiation phase. We observed highly dynamic auxin maxima during FM expansion, when *MpCYC* expressions remained dorsally restricted in WT but expanded broadly in *mpbop* FMs. These findings suggest that early symmetry breaking in the FM is guided by positional cues independent of auxin or BOP, which are instead essential for refining and maintaining dorsal-specific *CYC* expression during later FM enlargement. Our work illustrated how combining advanced imaging with emerging model systems can yield fresh insights into long-standing questions in development and evolution, and laid the groundwork for further elucidating the mechanisms underlying the repeated symmetry breaking in FMs.

## INTRODUCTION

Understanding how developmental novelties arise remains a fascinating challenge in biology, particularly when such novelties evolve repeatedly through the redeployment of the same molecular program across evolutionary scales. Bilateral symmetry (also called monosymmetry) in flowers is a striking representation of this phenomenon. Most flowers are either radially or bilaterally symmetric in their final forms ^1^. The former can be divided into equal halves along multiple axes, whereas the latter only exhibit a single plane of symmetry along the dorsal-ventral (D-V) axis, with dorsal side located closer to the inflorescence meristem (IM) than the ventral side (Fig. 1a). Monosymmetry has evolved at least 130 times independently from radial ancestors ^2^ and has been considered a key innovation in many species-rich lineages ^3,4^. Studies in snapdragon (*Antirrhinum majus*) in the early 1990s first revealed that two paralogous TCP transcription factors, *CYCLOIDEA* (*CYC*) and *DICHOTOMA*, were responsible for controlling dorsal organ identities in the flowers ^5–8^. Homologs of *CYC* have since been identified in numerous plant taxa, and strikingly, in almost all plant taxa examined to date, monosymmetry is associated with differential expression of *CYC* genes along the D-V axis ^9,10^.

**Figure 1.**
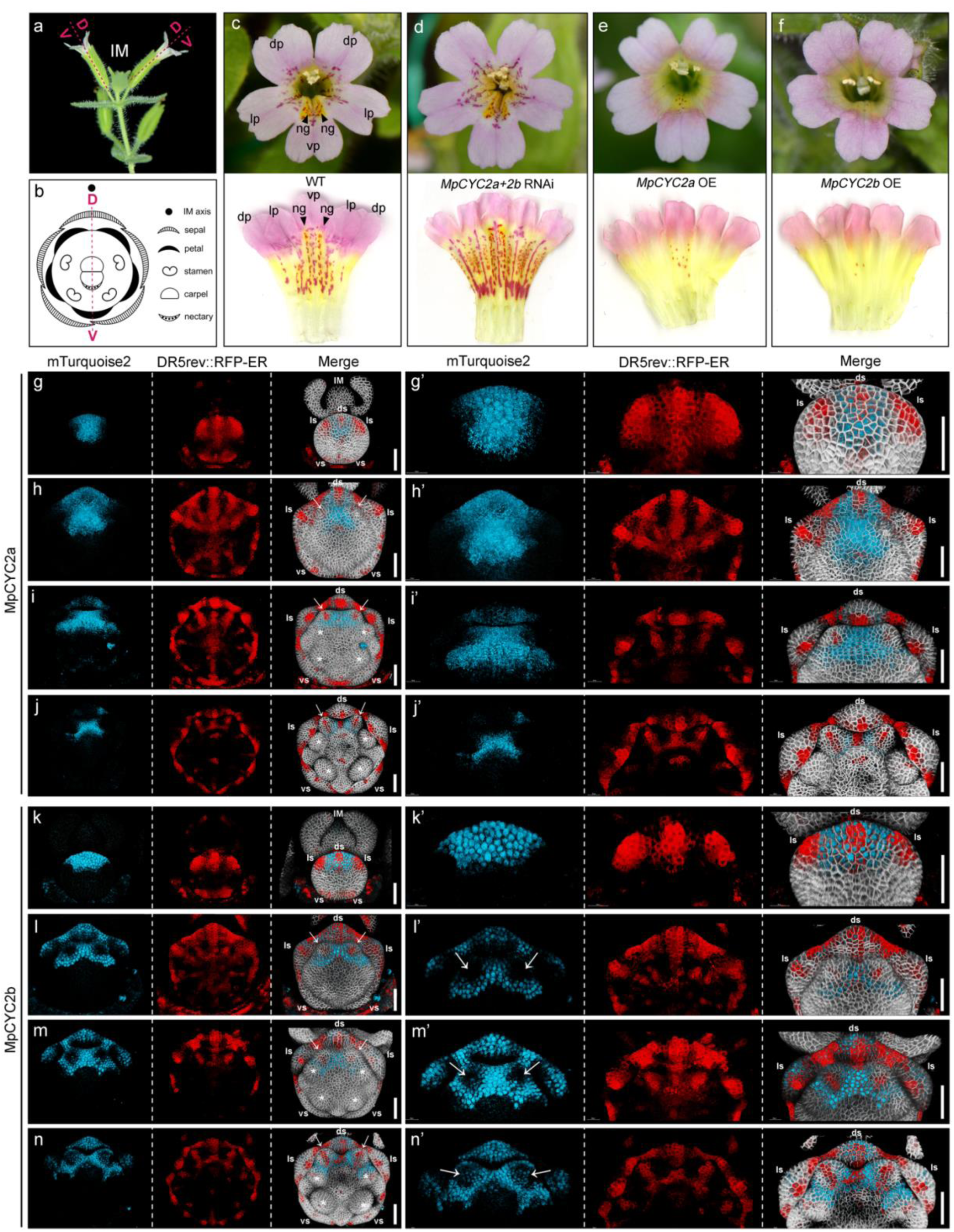
Mimulus parishii as a model for understanding mechanisms underlying floral bilateral symmetry. (a) Side views of *M. parishii* inflorescence. Dorsal sides of the flowers are located adjacent to the IM and the ventral sides are further away from the IM. (b) A floral diagram of *M. parishii* flower exhibiting bilateral symmetry. (c) A front view of a WT flower and its petal scan, black arrowheads pointing at the nectar guides on the ventral petals. (d) A front view of a *MpCYC2a/2b* double RNAi flower and its petal scans. (e) A front view of a *MpCYC2a* OE flower and its petal scans. (f) A front view of a *MpCYC2b* OE flower and its petal scans. (g-n) Maximum projections of MpCYC2a or MpCYC2b-mTurqoise2, DR5rev::RFP-ER, and merged with plasma membrane marker. Samples in g’-j’ are identical as those in a-h, but were zoomed in with a focus on the dorsal halves of the flowers. (g, g’, k, k’) Initiation of incipient sepal primordia. (h, h’, l, l’) Initiation of petal primordia. (i, i’, m, m’) Initiation of stamen primordia. (j, j’, n, n’) Initiation of carpel primordia. IM = inflorescence meristem; ds = dorsal sepal; ls = lateral sepal; vs = ventral sepal; dp = dorsal petals; ls = lateral petals; vp = ventral petals; asterisks in (g-n) = stamens; arrows in (g-n) = dorsal petals; ng = nectar guide; D = dorsal; V = ventral; All scale bars = 50 µm.

Despite three decades of research, how the *CYC*-based program is repeatedly recruited during evolution to generate bilaterally symmetric flowers remains unknown. This knowledge gap largely stems from the fact that in no plant system studied to date, the molecular mechanism establishing and maintaining the dorsal expression of *CYC* genes has been elucidated. Currently, only two potential regulators of *CYC* expression have been implicated. One is the phytohormone auxin. It was shown as early as 1961 that application of synthetic auxin could convert bilateral symmetry of *Tropaeolum* and *Linaria* flowers to ventralized radial symmetry ^11^. Similarly, exogenous applications of IAA (auxin), and sometimes NPA (auxin polar transport inhibitor), in snapdragon ^12^, *Matricaria inodora* ^13^, and *Mimulus lewisii* ^14,15^ were shown to trigger floral symmetry changes. Therefore, there is strong evidence that auxin can affect floral symmetry, but how such influence is implemented at the molecular level is unknown.

The other potential regulator of *CYC* expression is BLADE-ON-PETIOLE (BOP), a BTB-ankyrin protein that lacks DNA binding domains and acts as a transcription co-regulator, particularly as E3 ligase in ubiquitination complexes ^16^. *BOP* homologs appear to be highly pleiotropic and are involved in a wide range of developmental processes. Notably, *bop* mutants in *Arabidopsis, Medicago*, *Pisum*, *Hordeum*, and *Mimulus* all exhibit a change in flower symmetry ^17–21^. In bilaterally symmetric *M. lewisii* flowers, MlBOP has been shown to physically interact with MlCYC2a and MlCYC2b proteins; and *mlbop* mutant produces flowers with dorsalized characteristics due to ectopic expression of *MlCYCs* ^21^. But how are the interactions at protein level linked to the expansion of transcription domains remains puzzling.

One major obstacle impeding our understanding of bilateral symmetry establishment is that molecular mechanisms can be challenging to distill from genetic and phenotypic analyses alone. Transgenic manipulation and high-resolution imaging are often instrumental to link the dynamics of gene expression patterns to cellular processes that give rise to the phenotypic outcomes ^22–25^. Overcoming this obstacle requires systems with a functional monosymmetry program and equipped with suitable tools for molecular interrogations. However, most of the well-established models for studying flower development (e.g. *A. thaliana*, petunia, and tomato) produce radially symmetric flowers, and most plants with monosymmetric flowers (including the classic model snapdragon) remain inaccessible to routine stable transgenic manipulation.

Here, we developed the monkeyflower species *Mimulus parishii* as a model for mechanistic studies of floral symmetry establishment, for its naturally monosymmetric flowers and easiness in generating stable transgenic lines ^26–28^. By combining multiple fluorescent reporters and high-resolution confocal microscopy, we aimed to pinpoint when and where *CYC* expression is first initiated in the floral meristem (FM), and characterize the detailed expression dynamics during FM development, with a particular focus on pre-organogenesis stages. We captured incipient *CYC* expressions at the cellular resolution, shedding light on the positional information required for establishing D-V domains in the FM. Furthermore, we examined spatiotemporal patterns of *CYC* expression in relation to FM growth and auxin activity maxima in both wild-type (WT) and *bop* mutants. Our work unveiled distinct phases of *CYC* expression, clarified the timing when auxin and BOP act to maintain *CYC* expression in the dorsal half of the FM, and highlighted the complexity of the regulatory relationships among *CYC*, *BOP*, and auxin signaling. Together, these results serve as the crucial first step towards understanding the molecular mechanisms underlying repeated symmetry breaking in flowers.

## RESULTS

### MpCYC2a and MpCYC2b regulate floral symmetry in *Mimulus parishii*

To allow changes in floral symmetry easy to discern, we used an inbred line of *M. parishii* with much denser pigmentation spots on the ventral petal than the dorsal ones ^28^ (Fig. 1c, S1). Unlike dorsal and lateral petals that had flat surfaces, the ventral petal exhibited two slightly elevated ridges with yellow pigmentation and short hairs as nectar guides. In addition to petal morphology, other floral organs also display differences along the dorsal-ventral (D-V) axis that conferred bilateral symmetry (Fig. 1b, S1): dorsal sepals initiate earlier than other sepal primordia; the total number of stamens is four due to the missing dorsal stamen; the two lateral stamens are shorter than the two ventral stamens; and the nectary is only located at the base of the ventral carpel.

*MpCYC2a* and *MpCYC2b* are two *CYC* paralogs in *M. parishii*, resulted from the same Lamiale-specific duplication that gave rise to *CYC* and *DICH* in snapdragon ^29^. Similar to *CYC* and *DICH* in snapdragon, the two *CYC* paralogs had been shown to redundantly control dorsal organ identity in *M. guttatus* and *M. lewisii* ^21,30,31^. As expected, knocking down the expression of *MpCYC2a* and *MpCYC2b* (*MpCYCs*) in *M. parishii* via RNA interference (RNAi) led to ventralized flowers, while over-expression (OE) of either *MpCYC2a* or *MpCYC2b* resulted in flowers with dorsalized characteristics (Fig. 1d-f; Fig. S1). Both strong *MpCYC2a* and *MpCYC2b* OE lines showed additional phenotypes, including stunted vegetative growth, appearance of leaf pigmentation stripes and occasional ovule-to-carpel transformation (Fig. S1). In extreme cases, the vegetative part was exceptionally compact with very small leaves, and flowers would only produce on escaped shoots that resembled WT growth (Fig. S1).

### Dorsal expressions of MpCYC2a and MpCYC2b during organogenesis overlap but are not identical

To examine the expression patterns of MpCYCs and local auxin response at cellular resolution, we generated transgenic plants that contained 1) pUbi::RCI2A-mCitrine as a plasma membrane marker ^32^, 2) DR5rev::RFP-ER for translational readout of auxin activity maxima ^33^, and 3) translational reporters of MpCYC2a-mTurquoise2 and MpCYC2b*-*mTurquoise2 driven by their native promoters ^21^. To validate the efficacy of these reporter constructs, we first analyzed CYC expressions during organogenesis, starting from the emergence of sepal primordia until the initiation of carpel primordia (Fig. 1g-n), a developmental window where *CYC* expression patterns have been frequently examined by *in situ* hybridization in other plant species. Consistent with previous *in situ* hybridization results from various species ^5,29,34–37^, both MpCYCs showed dorsal-specific expression. However, our high-resolution imaging also revealed a few novel details.

For instance, while the amino acids sequences of both MpCYC2a and 2b have a nuclear localization signal and a predicted intrinsically disordered region (Fig. S2a, S2b), the nuclear localization was readily visible for only MpCYC2b but less evident for MpCYC2a; the latter formed numerous droplets that were dispersed in the cell (Fig. 1g-n, S2c). Furthermore, we saw a gradual reduction in the expression domains of MpCYC2a during organogenesis (Fig. 1g-j). At the incipient stage of sepal initiation, when DR5 signals clearly marked the five sepal initiation sites, expression of MpCYC2a covered almost half of the FM, from the dorsal-most region to the center of the FM (Fig. 1g, 1g’). When petal primordia started initiating, expression of MpCYC2a encompassed the entire dorsal sepal primordia and the region between the two dorsal petal primordia (Fig. 1h, 1h’). During the initiations of stamen and carpel primordia, expression of MpCYC2a in the dorsal sepal retreated and became restricted only between the two dorsal petals (Fig. 1i, 1i’, 1j, 1j’); this region corresponded to the missing dorsal stamen primordium.

On the other hand, such a reduction in expression domains as development progressing was not observed for MpCYC2b (Fig. 1k-n). During organogenesis, MpCYC2b expression remained confined to the dorsal and lateral sepals and the boundary regions surrounding the two dorsal petals (Fig. 1l-n, 1l’-n’). There appeared to be an exclusion of MpCYC2b proteins in initiating petal primordia (arrows in Fig. 1l, 1l’), but as petal development proceeded, MpCYC2b gradually extended into the petal primordia (arrows in Fig. 1m, 1m’, 1n, 1n’).

### Dorsal expressions of MpCYCs are established prior to any auxin maxima in the FM

We then tried to pinpoint when and where did MpCYC2a and 2b first emerge in the FM and examine how they relate to the spatiotemporal dynamics of auxin maxima using our triple reporter lines. While searching for the first appearance of MpCYC2a and 2b by dissecting FMs of very early stages (Fig. 2f), we encountered three surprising observations: Firstly, unlike *A. thaliana*, physical initiation of FMs in *M. parishii* was not accompanied by detectable auxin maxima in the FMs ^38^. Secondly, expressions of both MpCYCs were established prior to any DR5 signal in the FM. Thirdly, the initial activation of MpCYC2a and MpCYC2b occurred at different developmental stages and in different positions (Fig. 2, S3, S4).

**Figure 2.**
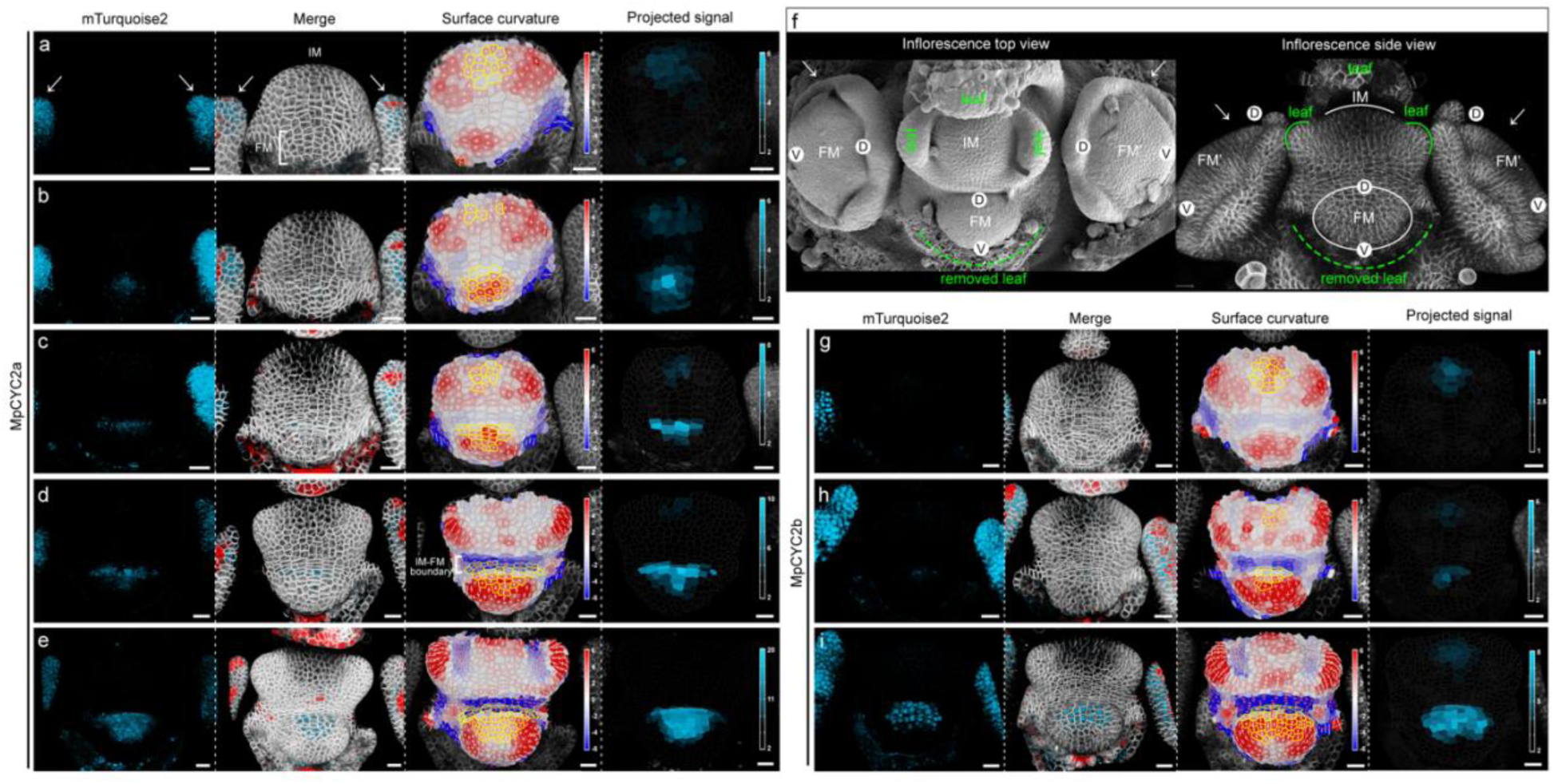
Initiation of MpCYC expressions in the FM. (a-e) Expression of MpCYC2a-mTurqoise2 during the earliest developmental stages of FMs. (f) Top view and side view of inflorescences of M. parishii. (g-i) Expression of MpCYC2b-mTurqoise2 during the earliest developmental stages of FMs. FMs in (c) and (g) are of similar developmental stages. Since no MpCYC2b expression was detected until (h), stages earlier than (g) were not shown. Columns in (a-e) and (g-i) from left to right were: maximum projection of MpCYC2a or 2b-mTurqoise2, maximum projection of all signals merged, surface Gaussian curvature, and heatmaps of relative intensity of mTurqoise2 signal projected onto the processed surface. For surface Gaussian curvature, cells had positive or negative curvatures were bulging out, or concaving inwards, respectively. Cells outlined yellow in the curvature panel are cells with a relative signal intensity above the 90^th^ percentile of all processed cells (i.e. all cells with a curvature value). Arrows in (a, f) were pointing at buds at the previous node. All scale bars = 20 µm.

In particular, the onset of MpCYC2a expression seemed to accompany the inception of FM. At the earliest stage of FM development that we could capture (Fig. 2a, S3a), although there was no discernable structure of the FM yet, a few cells in the region where FM initiation would take place exhibited weak but positive Gaussian surface curvatures (i.e. cells bulging out). Strong MpCYC2a-mTurquoise2 and DR5 signals were shown in the two floral buds at the previous node (arrows in Fig. 2a point to the dorsal sepal primordia from FMs of the previous node; see the two FM’ in 2f for examples), but they were not detectable in the incipient FM. Interestingly, a few cells in the IM displayed weak mTurquoise2 signal (Fig. 2a; cells outlined in yellow). The first appearance of MpCYC2a in the FM was detected when the number of cells in the incipient FM had a slight increase (Fig. 2b, S3b). At this stage, mTurquoise2 signals were found in almost all the cells in the incipient FM, in the IM, as well as few cells adjacent to the dorsal end of the FM that did not have positive surface curvature (i.e. not bulging out) (Fig. 2b). Subsequently, the physical initiation of FM became evident, so did the boundary between the IM and FM, which was marked by cells above the FM with increased aspect ratios and stronger negative curvatures (i.e. cells concave in). Although faint expressions of MpCYC2a-mTurquoise2 were detected in the IM throughout the early stages, signals in the FM significantly increased as the FM grew out, and cells with the highest expression levels became limited to those at the dorsal-most region of the FM and the boundary cells adjacent to it (Fig. 2c-e, S3c-e).

On the other hand, expression of MpCYC2b in the FM occurred later than that of MpCYC2a. Even when the expression of MpCYC2a had already became dorsalized in the FM (Fig. 2c), no signal of MpCYC2b was detected in the FM (Fig. 2g, which was of a comparable stage to 2c). When the IM-FM boundary cells became visibly concave, presence of MpCYC2b was first captured in just a few cells that are either the dorsal-most cells of the FM or boundary cells that are adjacent to them (Fig. 2h, S4b, S4d, S4e). As the development of the FM proceeded, the number of dorsal cells expressing MpCYC2b in the FM quickly increased (Fig. 2i, S4c). Similar to MpCYC2a, strong expression of MpCYC2b was also seen in the boundary cells at the dorsal end of the FM, and faint expression could be detected in a few cells in the IM.

### Highly dynamic auxin response pattern during FM expansion demarcates the ventral boundary of MpCYC expressions

We next investigated the spatiotemporal relationships between auxin maxima and MpCYCs during FM expansion, from when the FM just gained the dome-shape and became separated from the IM by the visibly distinguishable boundary cells (e.g. Fig. 2d, 2h) to the onset of organogenesis (i.e., the first sepal primordium became visible). During this developmental window, the FMs increased in size but maintained the typical dome-shape. However, patterns of auxin maxima appeared to be highly dynamic (Fig. 3a-f). DR5 signals were first observed when the FMs grew larger than 90 µm in width and 60 µm in height (Fig. 3a, 3f), in two small groups of cells that were located at the two ends of the mediolateral axis, followed by more DR5 cells appearing at the center of the FM (Fig. 3b). As the number of cells with DR5 signal increased in the center of the FM (Fig. 3b, 3b’), domain of expression started shifting towards the dorsal end of the FM (Fig. 3c). Subsequently, DR5 signals started to appear in cells at the ventral periphery of the FM (Fig. 3d). Next, the number of cells expressing DR5 dramatically increased in both dorsal and ventral domains (Fig. 3d, 3d’), but the band of cells along the mediolateral axis appeared to be depleted of DR5 signals (Fig. 3d’’). The number of cells expressing DR5 would then decrease, resolving into five patches that mark the initiation sites of sepals, followed by the dorsal sepal primordium emerging physically (yellow arrowhead in Fig. 3e).

**Figure 3.**
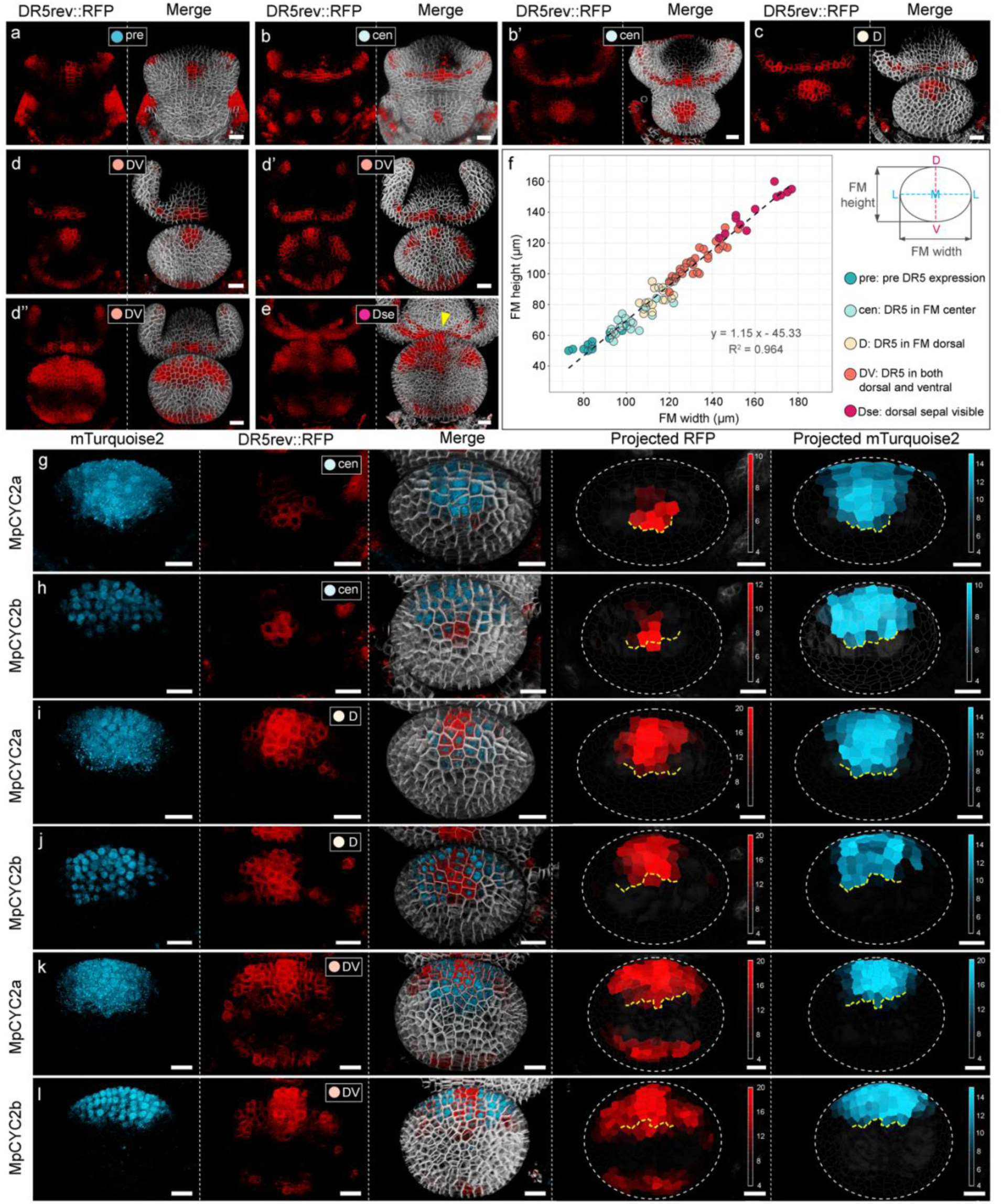
Auxin dynamics and MpCYC expression patterns during FM expansion. (a-e) Maximum projections of DR5rev::RFP-ER (left column) and merged with plasma membrane marker (right column) showing different DR5 patterns during early FM development. (f) Scatterplot of widths and heights of different FMs with their DR5 patterns color coded. (g-l) Expression patterns of MpCYC2a (g, I, k) and MpCYC2b (h, j, l) in FMs exhibiting “cen” (g, h), “D” (I, j), and “DV” (k, l) patterns of DR5. Columns from left to right were maximum projections of mTurqoise2, maximum projections of DR5rev::RFP-ER, merged maximum projections with plasma membrane marker, heatmap of DR5 signal of the sample, and heatmap of mTurqoise2 signal of the sample. Yellow dashed lines in the last two panels of (g-l) indicate the ventral-most boundary of MpCYC-mTurqoise2 expression boundaries. Yellow arrowhead in (e) pointing at a sepal primordium that had physically initiated. Scale bars = 20 µm.

The expressions of MpCYC2a and MpCYC2b remained dorsalized and largely overlapping during FM expansion (Fig. 3g-l). Intriguingly, the central or dorsal expression of DR5 seemed to define the ventral-most expression boundaries for MpCYC2a and MpCYC2b along the D-V axis during all stages (Fig. 3g-l; marked by the dashed yellow lines). We also examined DR5 patterns in *MpCYC2a* OE, *MpCYC2b* OE, and *MpCYC2a&2b* double RNAi lines. Surprisingly, no discernible differences in DR5 dynamics were observed in either of the OE backgrounds (Fig. S5). Meanwhile, in the double RNAi lines, the first appearance of DR5 expression was dorsalized, and no DR5 was detected in the FM prior to that (Fig. S6h). In other words, in the RNAi lines we did not observe the “cen” stage when cells in the center of the FM expressing DR5 (Fig. S6a vs. S6g) before the signal became dorsalized in the WT. Once cells on the ventral side also started to express DR5, the overall DR5 signal became somewhat disorganized (Fig. S6c-e vs. S6i-k). By the end of organogenesis, auxin response patterns seemed to become canalized again like the WT (Fig. S6f vs. S6l).

### Expression domains of MpCYC extended to ventral regions in *mpbop*

A recent study ^21^ showed that in *M. lewisii*, a species very closely related to *M. parishii*, both *MlCYC2a* and *MlCYC2b* were expressed ectopically with increased transcript levels in *mlbop* mutant, but questions remained as to when, where, and how MlBOP regulated their expression domains and levels. To address these questions, we designed CRISPR-Cas9 guides for *MpBOP* and obtained three independent *mpbop* lines (Fig. 4a). All mutant lines displayed the same set of phenotypes, including formation of extra lateral shoots, failure of organ abscission, occasional fasciation of the inflorescence stems and disruption of phyllotaxy, changes in floral organ numbers and identities, as well as tissue outgrowth on the inner side of sepals (Fig. 4b, S7). In particular, most flowers exhibited floral organ number irregularities, including fewer stamens and more sepals and petals (Fig. 4b, 4c). Similar to what was shown in *M. lewisii*, we observed the lateral and ventral petals of the majority of flowers acquiring dorsal petal morphologies (Fig. 4b), although the loss of ventral petal characteristics was rarely complete (Fig. S7). We then crossed *mpbop* with plants containing the triple fluorescent reporters to analyze changes in MpCYC expressions and auxin response patterns.

**Figure 4.**
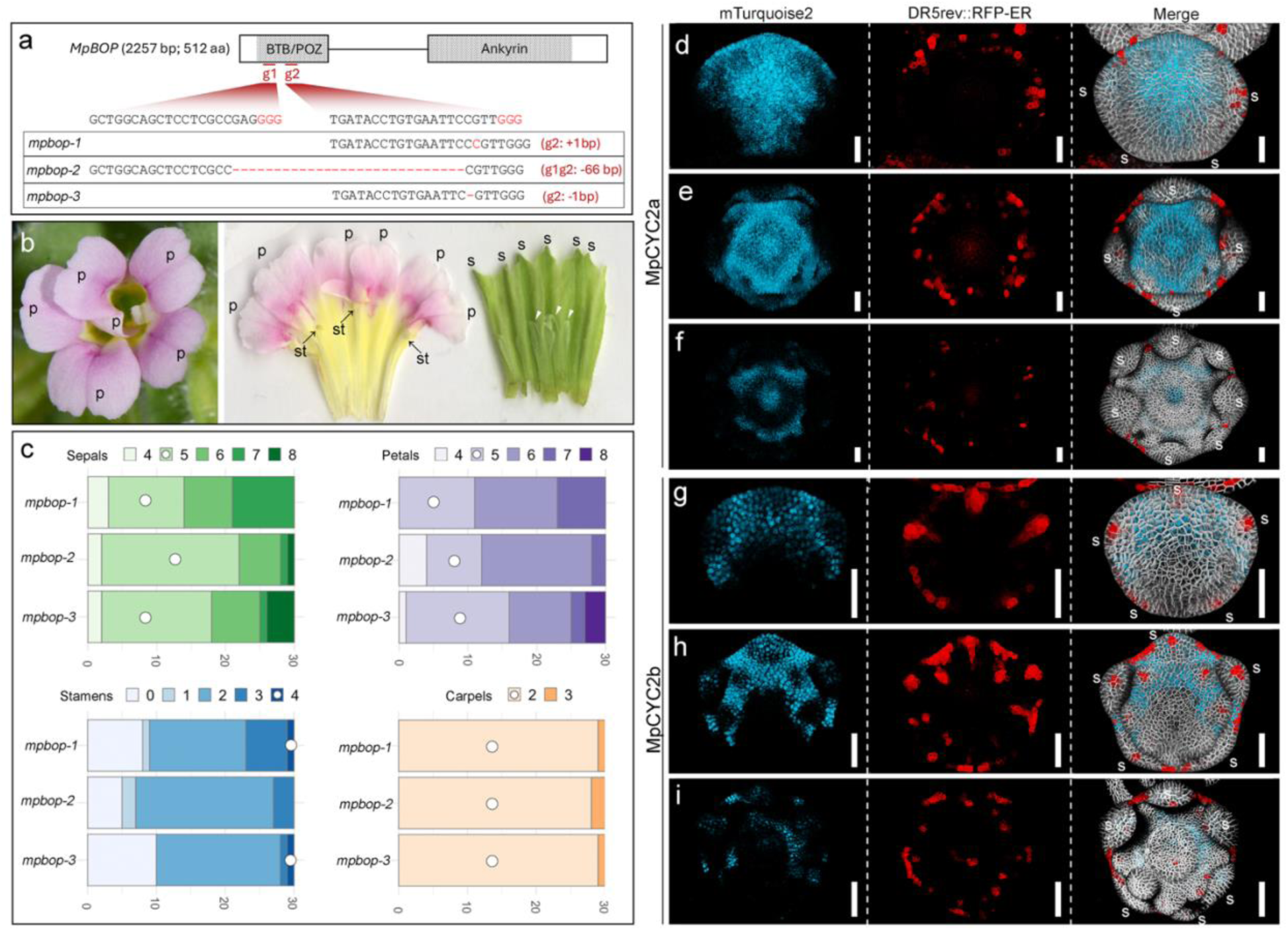
MpCYCs were ectopically expressed in mpbop. (a) CRISPR-Cas9 lines generated for *mpbop*. Three homozygous lines and their respective edits were shown. (b) Front view (left) and scan (right) of two *mpbop* flowers showing some representative phenotypes, including an increase petal and sepal numbers, a decrease in stamen numbers (arrows), and tissue outgrowth (white arrowheads) at the inner side of sepals. (c) Organ number counts of 30 flowers for each homozygous *mpbop* lines. Numbers with white circles were the organ numbers of a WT flower. (d-f) Patterns of MpCYC2a and DR5 in *mpbop*. (g-h) Patterns of MpCYC2b and DR5 in *mpbop* during organogenesis. Columns from left to right were maximum projections of MpCYC2a or 2b-mTurquoise2, DR5rev::RFP, and merged channels with plasma membrane marker. (d, g) A young flower bud with sepal primordia initiating. (e, h) A young flower bud with petal primordia initiating. (f, i) A young flower bud with all organs initiated. s = sepals; p = petals; st = stamens. Scale bars = 100 µm.

Consistent with the previous study ^21^, both MpCYC2a and 2b were ectopically expressed in *mpbop* flowers. However, the patterns of mis-expressions differed between the two paralogs (Fig. 4d-i). For instance, the expression of MpCYC2a extended to the ventral end along the D-V axis during sepal initiation (Fig. 4d), but became concentrated in between-whorl boundaries and the carpels during later organogenesis (Fig. 4e, 4f). On the other hand, although MpCYC2b expression also expanded in all stages examined, the extension was more prominent on the lateral sides than the center, and it had never reached the ventral-most regions along the D-V axis (Fig. 4g-i). Similar to what was observed in the WT, MpCYC2b also appeared to be excluded in the primordia initiation sites in *mpbop* (e.g. Fig. 4h).

### Initiation of MpCYC expressions was not affected in *mpbop*

We then investigated whether the initiation of MpCYC expressions was altered in the *mpbop* mutant. Expression of MpCYC2a could be detected in the IM at the earliest stages, also in almost all the cells in the incipient FM (Fig. 5a, 5b). Soon after FM initiation, MpCYC2a expression domain became dorsalized (Fig. 5c). Likewise, the expression of MpCYC2b was detected later than that of MpCYC2a (e.g. no expression in FM of Fig. 5d), and first appeared in a few cells located at the dorsal-most part of the FM and the adjacent IM-FM boundary region (Fig. 5e, 5e’). These observations demonstrated that the initiation of MpCYC expressions was not affected in *mpbop*.

**Figure 5.**
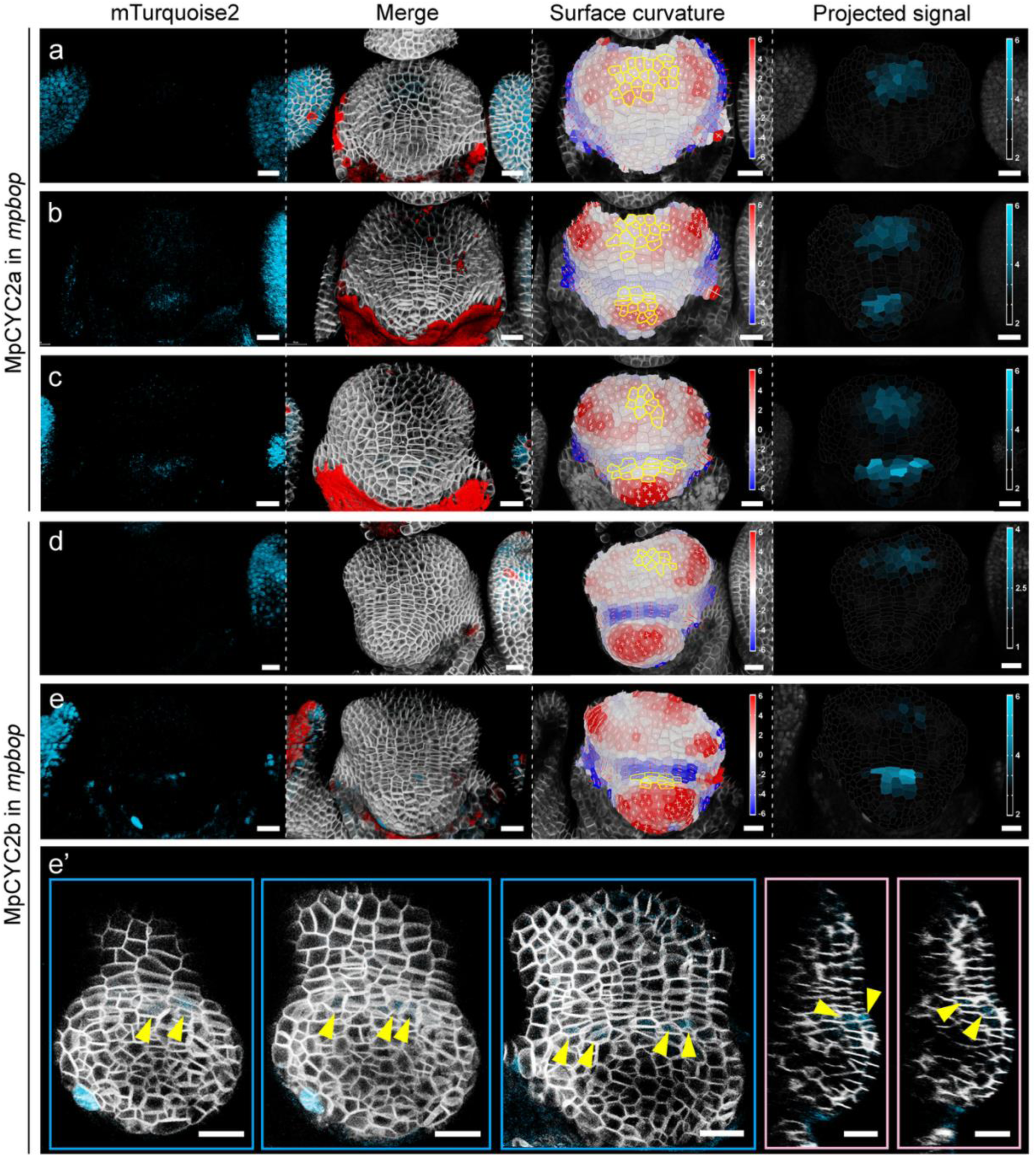
The spatiotemporal information of MpCYC activations remained unchanged in mpbop. (a-c) Expression of MpCYC2a-mTurqoise2 during the earliest developmental stages of *mpbop* FMs. (d, e) Expression of MpCYC2b-mTurqoise2 during the earliest developmental stages of *mpbop* FMs. Columns from left to right were: maximum projection of MpCYC2a or 2b-mTurqoise2, maximum projection of all signals merged, surface Gaussian curvature, and heatmaps of relative intensity of mTurqoise2 signal projected onto the processed surface. Cells outlined yellow in the curvature panel are cells with a relative signal intensity above the 90^th^ percentile of all processed cells (i.e. all cells with a curvature value). (e’) Z-stack slices (framed in blue) and orthogonal sections (framed in pink) through the center of the FM in (e) showing the cells expressing MpCYC2b. All scale bars = 20 µm.

### Ectopic expressions of MpCYCs coincided with onset of DR5 expression in the FM

Since the expression patterns of MpCYC2a and MpCYC2b remained unchanged in *mpbop* compared to the WT at the earliest stages (Fig. 5), yet ectopic expressions were observed already at the beginning of organogenesis (Fig. 4d-e), we sought to identify when the expression patterns started to diverge. To obtain this information, we first needed to examine whether the FM growth pattern differed between the WT and *mpbop*. Given the lack of obvious physical landmark for pre-organogenesis FMs, we characterized early FM development in WT and *mpbop* by combining FM size measurements and DR5 patterns.

In *mpbop*, the initial pattern of auxin maxima was similar to that of WT: no DR5 was observed at the earliest stages (Fig. 6a), and then two groups of DR5-expressing cells near the two ends of the mediolateral axis (Fig. 6b, S8) would appear. Another small group of cells with DR5 would then appear near the dorsal end of the FM (Fig. 6b’, S8). Subsequently, DR5 signals could be observed in a few clusters of cells along the ventral peripheries as the FM developed (Fig. 6c, 6c’, S8), followed by the physical emergence of sepal primordia (yellow arrowhead in Fig. 6d, S8). There was slight variation in the number and the relative locations of DR5-expressing cells among *mpbop* FMs at each stage (Fig. S8), but the general pattern that auxin maxima were first absent in the ventral side and appeared later was consistent. During these early developmental stages, the overall dimensions of the FM were very similar between the WT and *mpbop* (Fig. 6e). When aligning the DR5 dynamics in the WT to those in *mpbop*, it appeared that the onset of DR5 expression, first appearance of DR5 in the ventral side, and the initiation of sepal primordia, all occurred in FMs of similar sizes (Fig. 6f, 6h).

**Figure 6.**
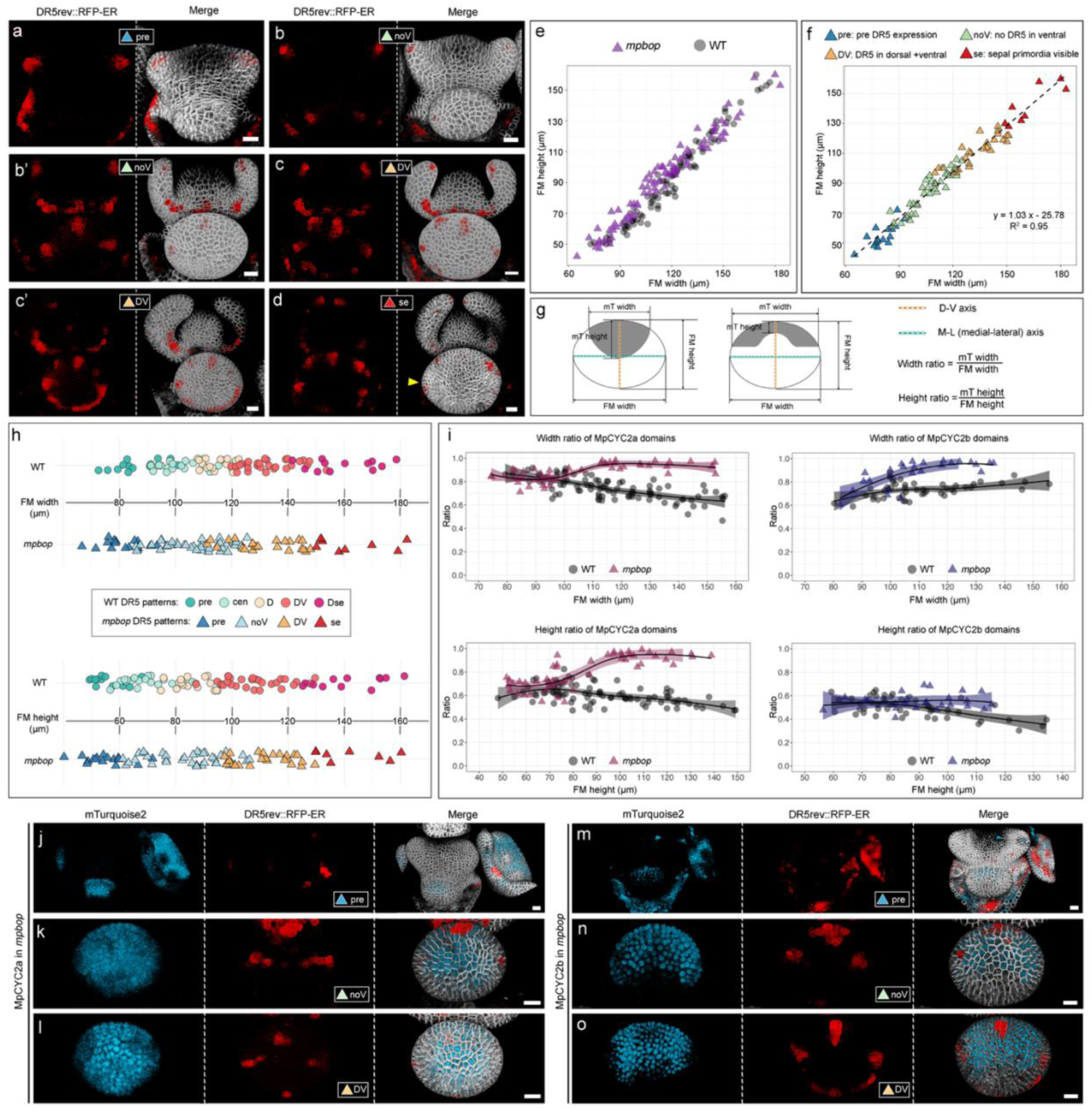
Domains of MpCYCs diverged in mpbop from WT coincided with onset of DR5 expression. (a-d) Maximum projections of DR5rev::RFP-ER (left column) and merged with plasma membrane marker (right column) showing different DR5 patterns during early FM development in *mpbop*. (e) Scatterplot of FMs of *mpbop* and WT from pre-DR5 to sepal primordia physically initiate stages. (f) Scatterplot of *mpbop* FMs with their DR5 patterns color coded. (g) Illustrated measurements of height and width ratios for MpCYC2a and MpCYC2b domains. (h) DR5 patterns of *mpbop* and WT FMs aligned by FM width (upper panel) and FM height (lower panel). (i) Measurement of FM width and height ratios of MpCYC2a and MpCYC2b domains in *mpbop* and WT FMs. (j-o) Expressions of MpCYC2a (j-l) and MpCYC2b (m-o) during early FM morphogenesis in *mpbop*. Columns from left to right were maximum projections of mTurqoise2, DR5rev::RFP-ER, and merged with plasma membrane markers, respectively. Yellow arrowhead in (d) pointing at a sepal primordium that had physically initiated. Scale bars = 20 µm.

Subsequently, we quantified the domains of MpCYC2a and MpCYC2b expression relative to the sizes of the FMs by measuring their height and width ratios (Fig. 6g): the former was calculated by using the length of the D-V axis of the FM and the length of MpCYC2a or MpCYC2b expression domain on that axis; the latter was calculated by using the width of the FM along the mediolateral axis and the width of the widest region of mTurqouise2 along the mediolateral axis. For MpCYC2a, both height and width ratios of the expression domains were initially almost identical in the WT and *mpbop*, but then both ratios started to quickly approach 1 in *mpbop*, while those of the WT showed a slight decline during the later developmental stages (Fig. 6i). The divergences of the height and width ratios for MpCYC2a occurred when the FM reached about 100 µm in width or 70 µm in height, coinciding with the size range showing the onset of DR5 expression for both the WT and *mpbop* (Fig. 6h). In other words, the expression domains of MpCYC2a were not significantly different in WT and *mpbop* before the occurrence of auxin maxima (Fig. 6j), after which the domains rapidly expanded both laterally and along the D-V axis in *mpbop* (Fig. 6k, 6l). Similarly, MpCYC2b expression was nearly identical between the WT and *mpbop* prior to visible DR5 signal (Fig. 6i, 6m), and the divergence of width ratios occurred when the FM reached ∼90 µm in width. Interestingly, the height ratios of MpCYC2b stayed very similar in those two backgrounds (Fig. 6i). Consistent with what was observed during organogenesis (e.g. Fig. 6g-i), the expression domains of MpCYC2b during FM expansion extended mostly laterally in *mpbop*, but not along the D-V axis (Fig. 6n, 6o).

## DISCUSSION

In this study, we combined transgenic manipulation and confocal microscopy to investigate the spatiotemporal details of symmetry breaking in the FMs of *M. parishii*. Our cellular-resolution imaging of multiple fluorescent reporters enabled us to directly visualize the expression dynamics of MpCYC2a and MpCYC2b, two key genes controlling monosymmetry. By correlating MpCYC expressions with patterns of auxin maxima and FM growth in both wild type and the *mpbop* mutant, we could recognize three phases of MpCYC expressions (Fig. 7): (I) Initiation. This is the phase before the FM reaches ∼100 µm in width, with no detectable DR5 signal in the FM. By the end of this phase, both MpCYC2a and 2b have fully established dorsal expression domains in the FM (Fig. 2e and 2i); (II) Maintenance. This is the phase between the emergence of the first DR5 signal and the onset of organogenesis (FM reaching ∼150 µm), during which MpCYC2 expressions are maintained in the dorsal half of the wild-type FM (Fig. 3g-3l) but become massively expanded in the *mpbop* FM; (III) Specification. This phase coincides with organogenesis, during which MpCYC expressions gradually become restricted to specific domains associated with dorsal organs (Fig. 1g-n).

**Figure 7.**
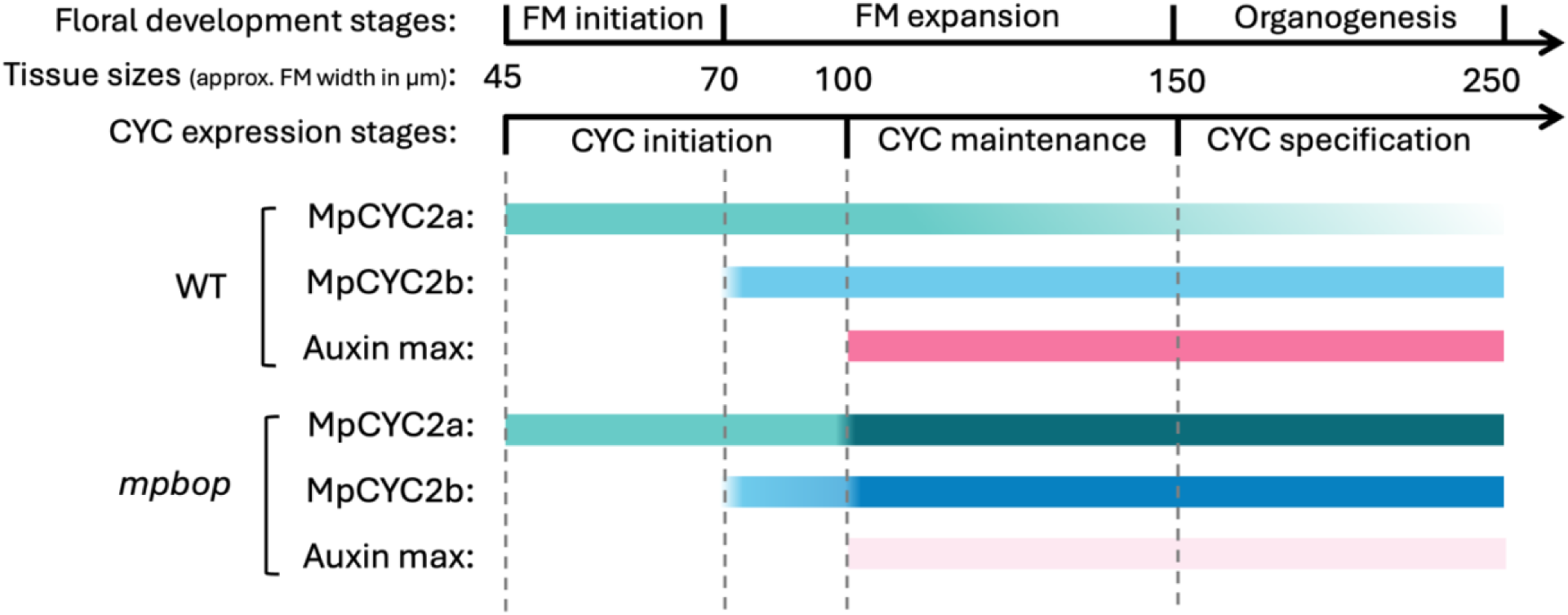
Distinct phases of MpCYC expressions during early floral development. Diagram showing distinct phases of CYC expressions in relationship to early floral developmental stages represented by approximate FM width. Relative expression levels of MpCYCs and DR5 were represented by the intensity of the colors; lighter and darker colors indicated lower or stronger expression levels, respectively.

Recognizing these distinct phases allowed us to clarify the timing when auxin or BOP affects MpCYC expressions. Our results suggest that neither auxin nor MpBOP is involved in the regulation of MpCYC expressions during the initiation phase. Both MpCYC paralogs already gained the dorsal specific expression in the FM prior to the first detectable auxin maxima, and their expression patterns remained unchanged in the *mpbop* mutant until the FM reached ∼100 µm in width, a stage coinciding with the onset of DR5 expression during FM expansion (Fig. 6). Together, these observations suggest that auxin and MpBOP most likely play key roles in maintaining, rather than initiating, MpCYC expressions. A similar phenomenon, where the initiation and maintenance of the same developmental program are controlled by different molecular modules, has been documented in adaxial-abaxial polarity establishment in leaves ^22,39^.

Although our results provide strong evidence that both auxin and MpBOP are involved in maintaining the dorsal specific expression of MpCYCs, the data also highlight the complexity of their regulatory relationships. Exogenous hormone treatments in previous studies^11–15^ implied that auxin could inhibit *CYC* expression. Consistently, our imaging data revealed negative correlations between DR5 and MpCYC signals in some cells (e.g., at petal primordium initiation sites; Fig. 1), although these inhibitory effects appeared highly context dependent. On the other hand, DR5 expression patterns were also altered in the *MpCYC2a&2b* double RNAi lines (Fig. S6), indicating that MpCYCs can also feedback to auxin signaling.

Our detailed examination of auxin response dynamics in the FM also revealed some surprising differences from what is known in *A. thaliana*^22,40^. Firstly, physical initiation of the FM in *M. parishii* was not accompanied by auxin maxima (Fig. 2). Secondly, auxin response patterns in the FM were highly dynamic during FM enlargement (Fig. 3), and most remarkably, there is a consistent overlap between the ventral-most boundary of DR5 and MpCYC expression domains (Fig. 3, dashed yellow lines). This raises the possibility that an auxin-dependent pathway defines the ventral boundary of MpCYC expressions at these stages. It is important to note that DR5 patterns reflect the integration of all auxin-related pathways within a tissue. Delineating the precise relationship between auxin and *MpCYCs* will require identifying specific effector genes in the auxin signaling pathway, examining their spatiotemporal expression patterns, and analyzing how they influence *MpCYC* expressions across developmental phases.

In *M. lewisii,* MlBOP and MlCYCs proteins physically interact ^21^. Given that the DNA sequences of *MpBOP* and *MpCYCs* are nearly 98% identical to *their M. lewisii* orthologs, it is reasonable to assume that these physical interactions are conserved in *M. parishii*. However, the ectopic expressions of the MpCYCs observed in *mpbop* mutants indicated that MpBOP could also influence *MpCYCs* at transcriptional levels. This regulation is most likely through interaction with other co-factors, as BOP proteins lack DNA-binding domains. The nature of MpBOP regulation of MpCYCs could be better revealed by studying the expression dynamics of MpBOP at the same time. Unfortunately, we were not able to obtain a functional reporter for MpBOP after multiple attempts. One plausible scenario is that MpBOP regulates *MpCYC* transcriptions via auxin-dependent pathways. In *mpbop*, auxin response levels were generally reduced, and the stage of ectopic MpCYC expressions coincided with the onset of DR5 activity. Although the onset of DR5 expression seemed to be consistent temporally in the WT and *mpbop* (Fig. 6, 7), their spatial patterns differed, suggesting that auxin activity in specific FM cells, possibly mediated by MpBOP, was necessary to restrict MpCYC expressions to the dorsal domain.

Lastly, our results also provided a few important clues to the quest of finding the upstream regulators of *MpCYCs*. The faint but consistent signals of both genes in the IM suggest that positional cues may originate from the IM itself, supporting previous hypotheses that the IM and subtending bract provide spatial information to establish D–V polarity ^41^. Additionally, the early expression of both MpCYCs were observed in the boundary cells between the IM and FM, and particularly, the onset of MpCYC2b was always observed at the IM-FM boundary, suggesting that boundary factors might also contribute to their regulations.

Understanding gene function in development requires connecting spatiotemporal expression patterns with the local cellular processes that ultimately give rise to phenotypic outcomes. Without fluorescent reporter systems and high-resolution imaging, building such connections would be infeasible. Our knowledge of the CYC homologs exemplifies this challenge: despite decades of research establishing their pivotal role in governing bilateral symmetry in numerous species, the processes and mechanisms that determine their precise expression domains remained elusive. To this end, our current work uncovered previously unknown details regarding the three distinct phases of *CYC* expression in relation to auxin maxima and BOP activity, and laid the groundwork for further elucidating the molecular networks underlying the repeated symmetry breaking in the FM.

## MATERIALS AND METHODS

### Plant materials and growth conditions

The near-isogenic line of *M. parishii* was generated in ^28^. All seeds were germinated on wet soil (Lambert LM-GPS germination mix) and stratified at 4°C for at least a week. After germination, all plants were grown in the University of Connecticut Ecology and Evolutionary Biology (EEB) research greenhouses, under natural light supplemented with vapor lamps to ensure a 16-hour light and 8-hour dark photoperiod. Plants were fertilized twice per week.

### Construction of transgenic lines

Plasmids for plasma membrane marker *pUBQ*::mCitrine-RCI2A and auxin reporter DR5rev::RFP-ER were previously published in ^32^ and ^33^, respectively. CDS of *MpCYC2a* and *MpCYC2b* were amplified and cloned into intermediate plasmids pDONR207 through BP reaction (Invitrogen, CA, USA). The respective intermediate plasmids were then cloned into pEarleyGate 101 through LR reaction (Invitrogen, CA, USA) to create the over-expression constructs for MpCYC2a and MpCYC2b. *MpCYC2a* and *2b* double RNAi lines was generated by amplifying a short coding DNA sequence of each gene from *M. parishii* cDNA, combining the sequences into one by PCR, and cloning into vector pB7GWIWGII ^42^ via Gateway cloning (Invitrogen, CA, USA).

To make transcriptional reporters of *MpCYC2a* and *MpCYC2b*, their respective 3kb promoters plus CDS were amplified via PCR into pENTR™/D-TOPO™ vector (Invitrogen, CA, USA), linearized, then cloning into pMCS-GW (Addgene #209360) via LR reaction. The CDS of mTurquoise2 was cloned from plasmid pSW646-pML1 (Addgene #115995) and incorporated in the final plasmids via NEBuilder® HiFi DNA Assembly (New England Biolabs, MA, USA).

The endogenous tRNA-processing CRISPR-Cas9 system was used to generate the generate *MpBOP CRISPR* lines. Guide spaces sequences were selected using CRISPR-P v2.0 (http://crispr.hzau.edu.cn/cgi-bin/CRISPR2/SCORE) and amplified using primers listed in Table S1. Amplified spacers were then incorporated into linearized backbone pRGEB32-BAR-AtU6.29 ^28^ using NEBuilder® HiFi DNA Assembly (New England Biolabs, MA, USA).

All final plasmids were verified by PCR and whole plasmid sequencing, and were transformed into Agrobacterium tumefaciens GV3101 following procedures in ^43^. *MpBOP* CRISPR lines were confirmed with PCR and sanger sequencing.

All primers are listed in Table S1.

### RNA extraction and qRT-PCR

For RNA preparation, 5 mm whole flower buds were collected and placed into liquid nitrogen immediately. Three biological replicates were collected for each sample type. Frozen plant tissue was ground using a Mini-G Geno/Grinder tissue homogenizer and 4 mm stainless steel grinding balls (Spex Sample Prep, Metuchen, NJ) for 2 minutes. RNA was extracted using a Spectrum Plant Total RNA Kit (Sigma-Aldrich, Saint Louis, MO) and eluted in a final volume of 40ul. Complementary DNA (cDNA) was synthesized from 500ng of RNA using the GoScript Reverse Transcription system (Promega, Madison, WI). Quality of the cDNA was checked by amplifying a large gene (5kb), *MpZEP2* indicating that the cDNA is intact and not degraded. RT-qPCR experiments were conducted using a CFX96 touch real-time PCR detection system (Bio-Rad, Hercules, CA) with SYBR Green master mix (Applied Biosystems, Foster City, CA). Primers used for qRT-PCR are listed in Table S1. The relative expression of the target genes was normalized by the expression of a reference gene *MpUBC*. Relative expression values were calculated using the delta-delta CT method. The statistical significance of RT-qPCR data was calculated using Welch’s T-tests with Microsoft Excel.

### Scanning Electron Microscopy

Tissues were dissected under a dissection microscope and fixed in FAA (50% ethanol, 10% formalin, 5% glacial acetic acid, and 35% water) overnight at room temperature with gentle agitation. The day before imaging, tissues were then gradually dehydrated through an ethanol gradient (50%, 75%, 85%, 95%, 100%) and stored in 100% ethanol overnight with gentle agitation. Samples were then critical point dried, sputter coated with gold-palladium, and imaged with FEI Nova NanoSEM 450 Scanning Electron Microscope at the UConn Bioscience Electron Microscopy Laboratory.

### Confocal Microscopy

For both single time imaging and live-imaging, tissues were dissected under a dissection scope and transferred to 55mm plates containing autoclaved 2% Sucrose, 1% Agarose, 0.5x Linsmaier and Skoog Medium, and 0.1% Plant Protective Medium (Plant Cell Technologies, DC, USA; added after autoclaving when the medium is cooled but not solidified). Samples were then immediately imaged on Leica SP8 Spectral Confocal at the UConn Advanced Light Microscopy Facility. For each fluorescent protein, the excitation (ex), emission (em), receptor and laser parameters were as following: for mTurquoise2, ex = 458 nm, em = 463-507 nm, HyD4 detector, gain 90%, laser power 20%; for mCitrine, ex = 514 nm, em = 519-700 nm, PMT2 detector, gain 700, offset -2%, laser power 10%; for RFP, ex = 561 nm, em = 604-658 nm, PMT5 detector, gain 800, offset -1%, laser power 5%. All images were acquired with 20x water immersion lenses, z-step size 1µm, collected in 16 bits and in 1024×1024 pixel resolution.

### Image analysis

Maximum projections of different fluorescent channels and the merged images were generated using LAS X Life Science software (Leica, Wetzlar, Germany). Orthogonal sections of samples were obtained using Fiji ImageJ after using the Transform-Rotate function to make sure the section can go through the center of the meristems.

Surface curvature and projected signals were processed using MorphographX following the manual ^44,45^. Briefly, surface detection was performed with the “Edge Detect” tool with a threshold from 4000-8000, depending on the brightness of the sample; Initial meshes were created with a 5 µm cube size and all meshes were smoothed and subdivided three times before projecting the membrane signal (1-2 µm from the mesh). Segmentation was performed with the “Auto-segmentation” tool, followed by manual corrections of segmentation errors. Surface Gaussian Curvature was computed with a radius of 20 µm. All mTurquoise2 signal projection were performed at the same 1-10 µm from the mesh and generated by running Mesh/Heat map/Measures/Signal/Signal Interior.

To outline cells with high mTurquoise2 signal onto the Gaussian curvature heatmap, after mTurquoise2 signals were projected onto the mesh, data of all processed cells and their respective signal levels were exported into a .cvs file using the Mesh/Attributes/Save to CSV function. The values of 90^th^ percentile of all processed cells were then calculated in Microsoft Excel. Subsequently, the .csv files were reloaded into the heat map, but cells with strong expression were selected using the Mesh/Heat map/Heat Map Select function, with the lower selection value as the calculated 90^th^ percentile value, and a higher value that is larger than the strongest signal value. Lastly, re-run the Gaussian curvature heatmap with the same radius of 20 µm.

## AUTHOR CONTRIBUTIONS

Y.M. and Y-W. Y. designed the study. B.F. constructed the RNAi lines, performed genotyping and qRT-PCR. All other experiments and analyses were performed by Y.M. The manuscript was written by Y.M. and revised by Y-W. Y.

## ACKNOWLEDGEMENT

We would like to thank Chang Liu, Matt Opel, and Meghan Moriarty for exemplary plant care at the UConn Botanical Conservatory. We appreciate comments from Yan Gong, Elena Kramer, and the Yuan lab members on improving the manuscript. This work was supported by NIH grant R01GM140092 (Y-W. Y.) and NSF award 2209220 (Y. M.). The Leica SP8 confocal microscope was acquired with a NIH shared instrumentation grant (S10OD016435) awarded to Akiko Nishiyama.

## SUPPLEMENTAL MATERIALS

**Figure S1.**
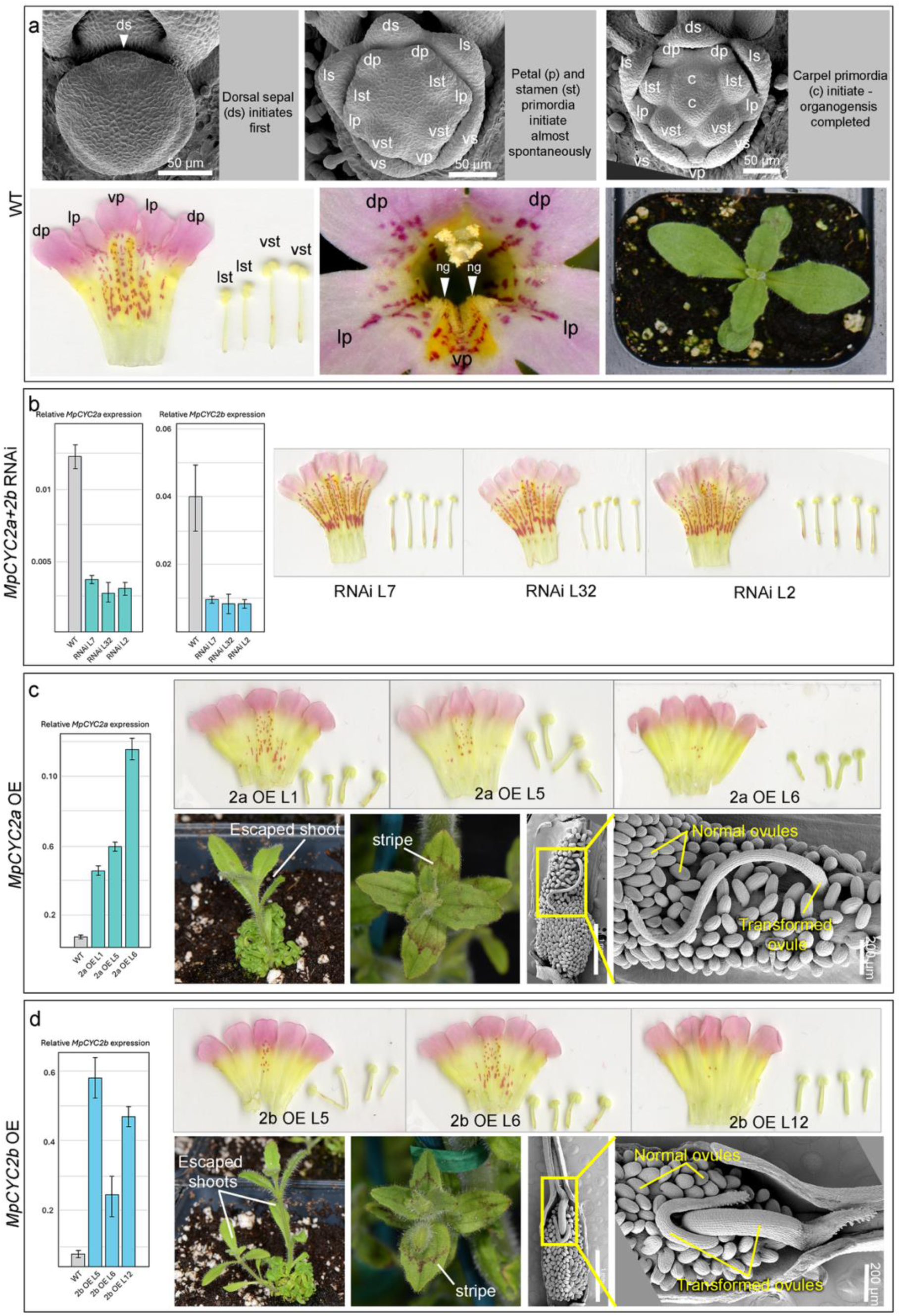
Mimulus parishii as a model for understanding mechanisms underlying floral bilateral symmetry. (a) WT *M. parishii*. Upper panels showed the ontogeny series of organogenesis in WT. Lower panels include petal and stamen scan of a WT flower (left), close up of a WT flower showing characters of nectar guides (ng; middle), and morphology of a WT seedling (right). (b) qRT-PCR (left) and petal and stamen scans (right) of three *MpCYC2a*+*MpCYC2b* double RNAi lines. All petals became ventralized and stamen numbers increased to five (c) qRT-PCR and phenotypes of *MpCYC2a* OE lines. Upper panel included petal and stamen scans of three OE lines. Lower panel included rare OE phenotypes of extremely compacted shoots, pigmentation stripes on leaves, and ovules transformed into carpels. (d) qRT-PCR and phenotypes of *MpCYC2b* OE lines. Upper panel included petal and stamen scans of three OE lines. Lower panel included rare OE phenotypes of extremely compacted shoots, pigmentation stripes on leaves, and ovules transformed into carpels.

**Figure S2.**
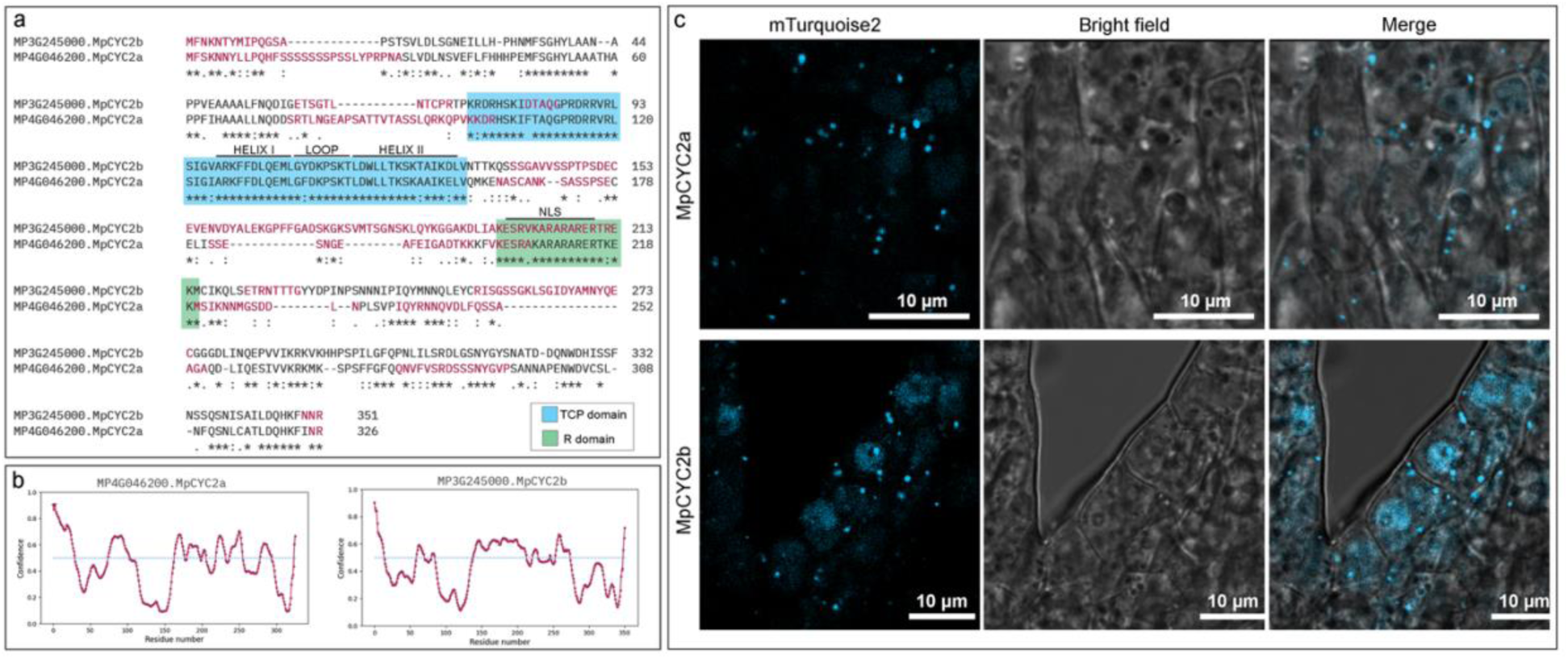
Amino acid sequence comparisons between MpCYC2a and MpCYC2b. (a) Amino acid alignment of MpCYC2a and MpCYC2b. Conserved domains and motifs were highlighted. Amino acids in red were predicted to have high likelihood of form intrinsically distorted regions shown in (b). (b) Screenshots of disorder profile plots of MpCYC2a and MpCYC2b from PrDOS (https://prdos.hgc.jp/cgi-bin/top.cgi). (c) Expression of MpCYC2a and MpCYC2b in older (∼ 5 mm) petals. Both show nuclear localized signals and condensates in the cytoplasm, but nuclear location of MpCYC2a was much weaker than that of MpCYC2b.

**Figure S3.**
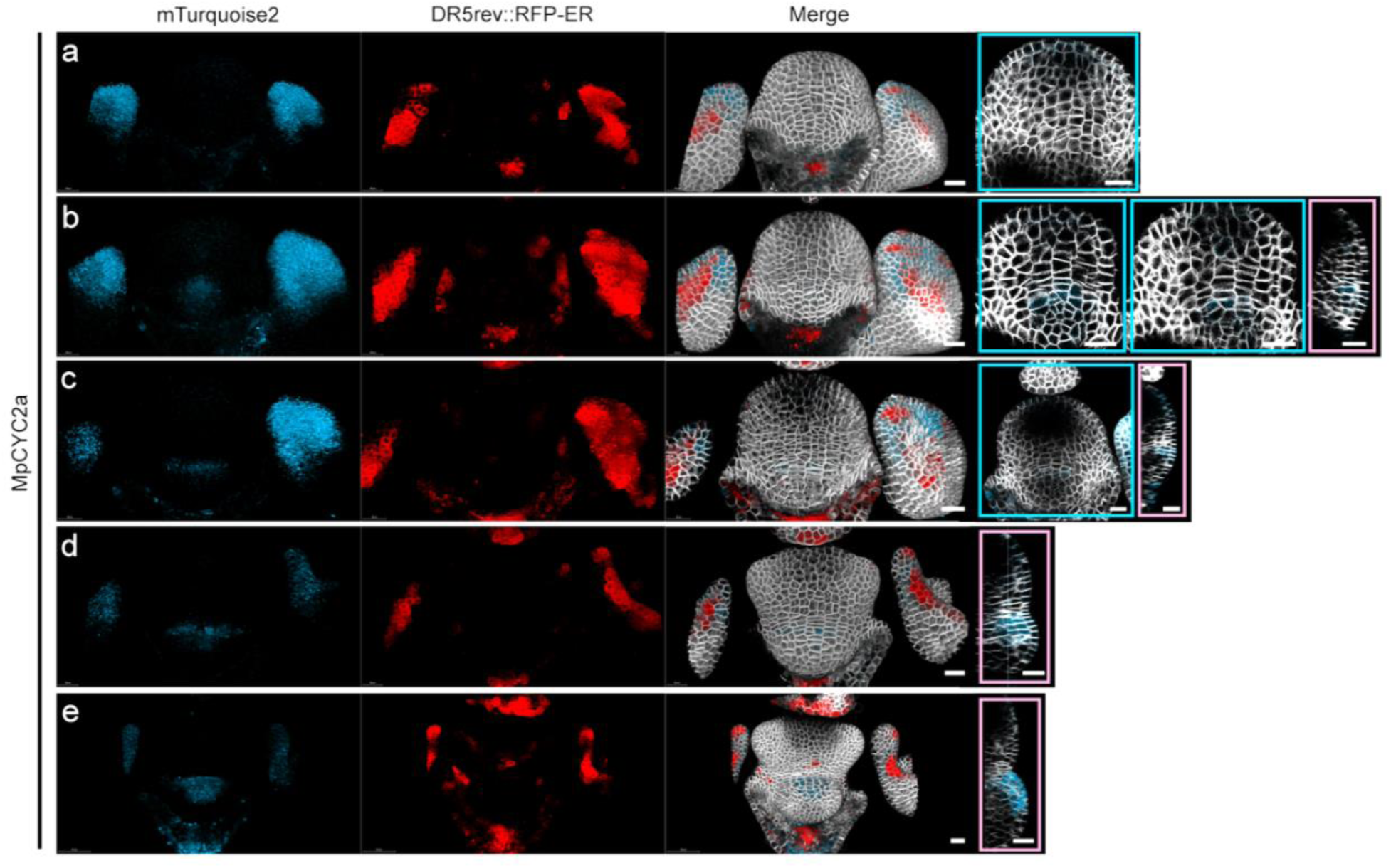
Initiation of MpCYC2a expression in the FM. (a-e) Original, uncropped versions of Fig. 2a-e, showing maximum projections of mTurqoise2 (first column), DR5rev::RFP-ER (second column), and merged with plasma membrane marker (third column). Images in blue frames showing merged z-stack slices and images in purple frames showing merged orthogonal sections through the center of the FM. All scale bars = 20 µm.

**Figure S4.**
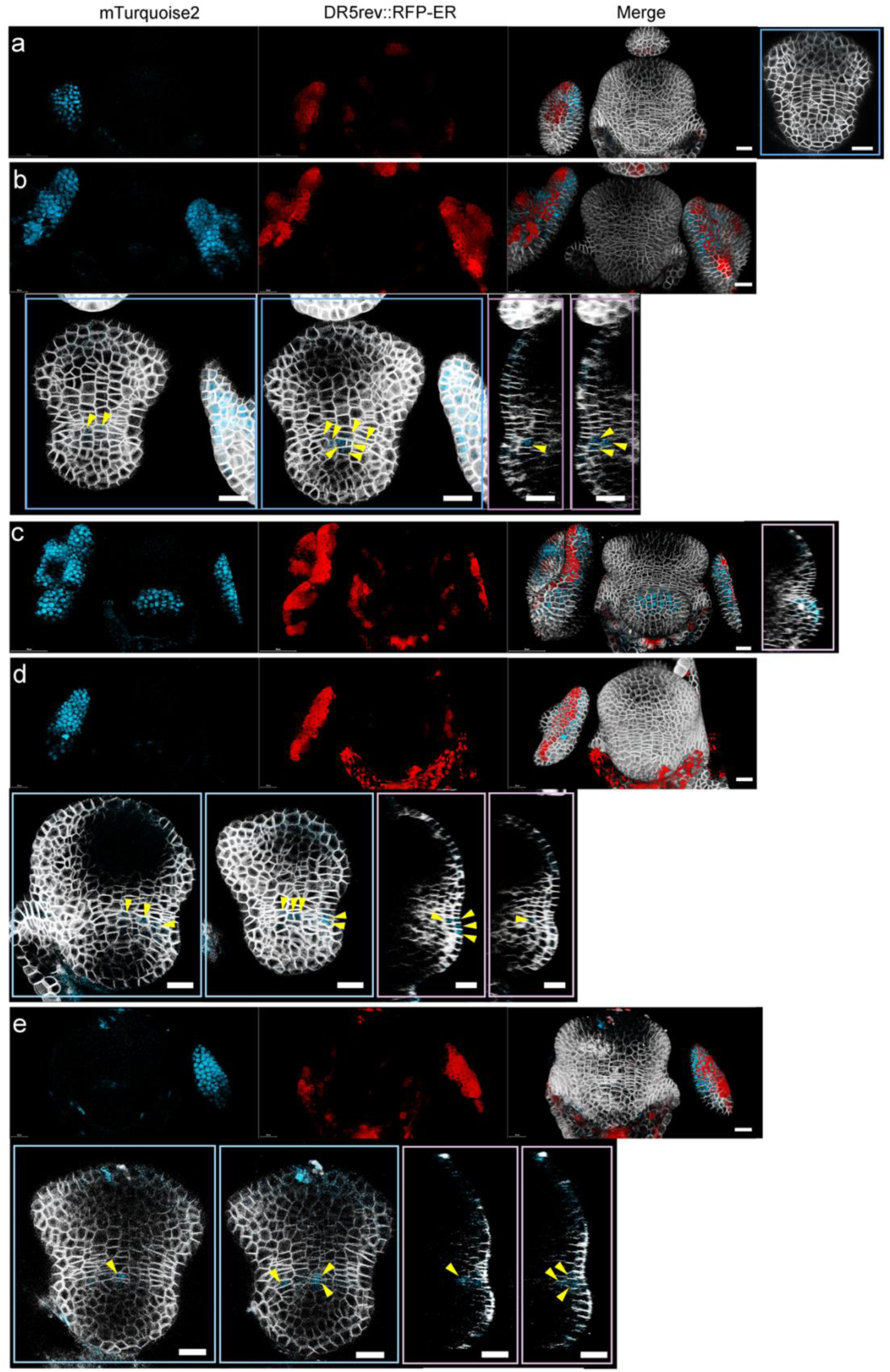
Initiation of MpCYC2b expression in the FM. (a-c) Original, uncropped versions of Fig. 2g-i, showing maximum projections of mTurqoise2 (first column), DR5rev::RFP-ER (second column), and merged with plasma membrane marker (third column). (d, e) Two additional samples with MpCYC2b-mTurqoise2 showing the earliest stages of MpCYC2b expression can be detected in just a few cells at the dorsal-most region of the FM and the adjacent IM-FM boundaries. Images in blue frames showing z-stack slices and images in purple frames showing orthogonal sections through the center of the FM. All scale bars = 20 µm.

**Figure S5.**
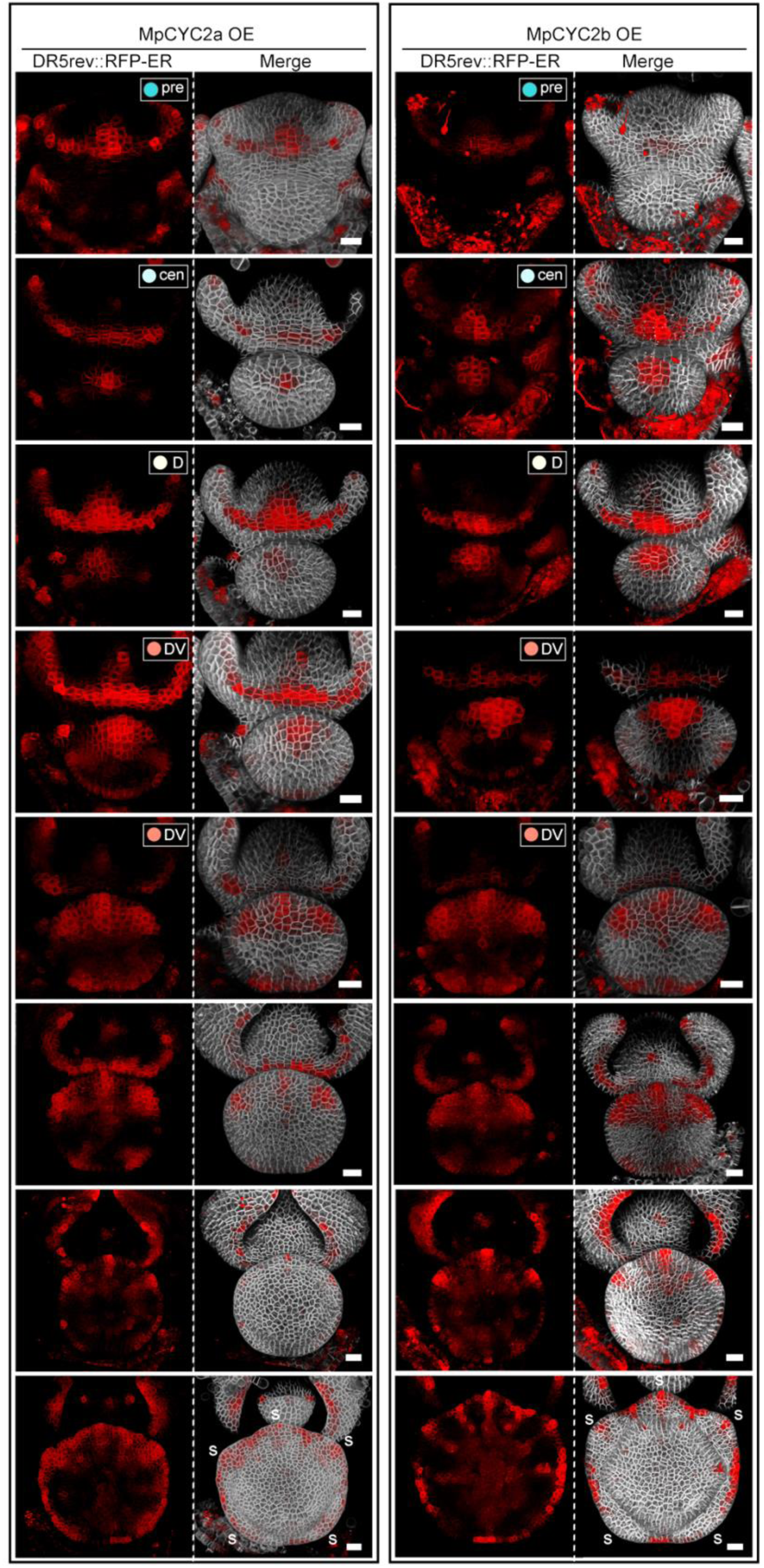
Early FM DR5 dynamics patterns did not change in MPCYC2a *OE nor MpCYC2b OE lines.* DR5 patterns in *MpCYC2a* OE (left) and *MpCYC2b* OE (right) during early FM morphogenesis and organogenesis. Equivalent FM DR5 stages (i.e. pre, cen, D, DV) were marked. s = sepals. Scale bars = 20 µm

**Figure S6.**
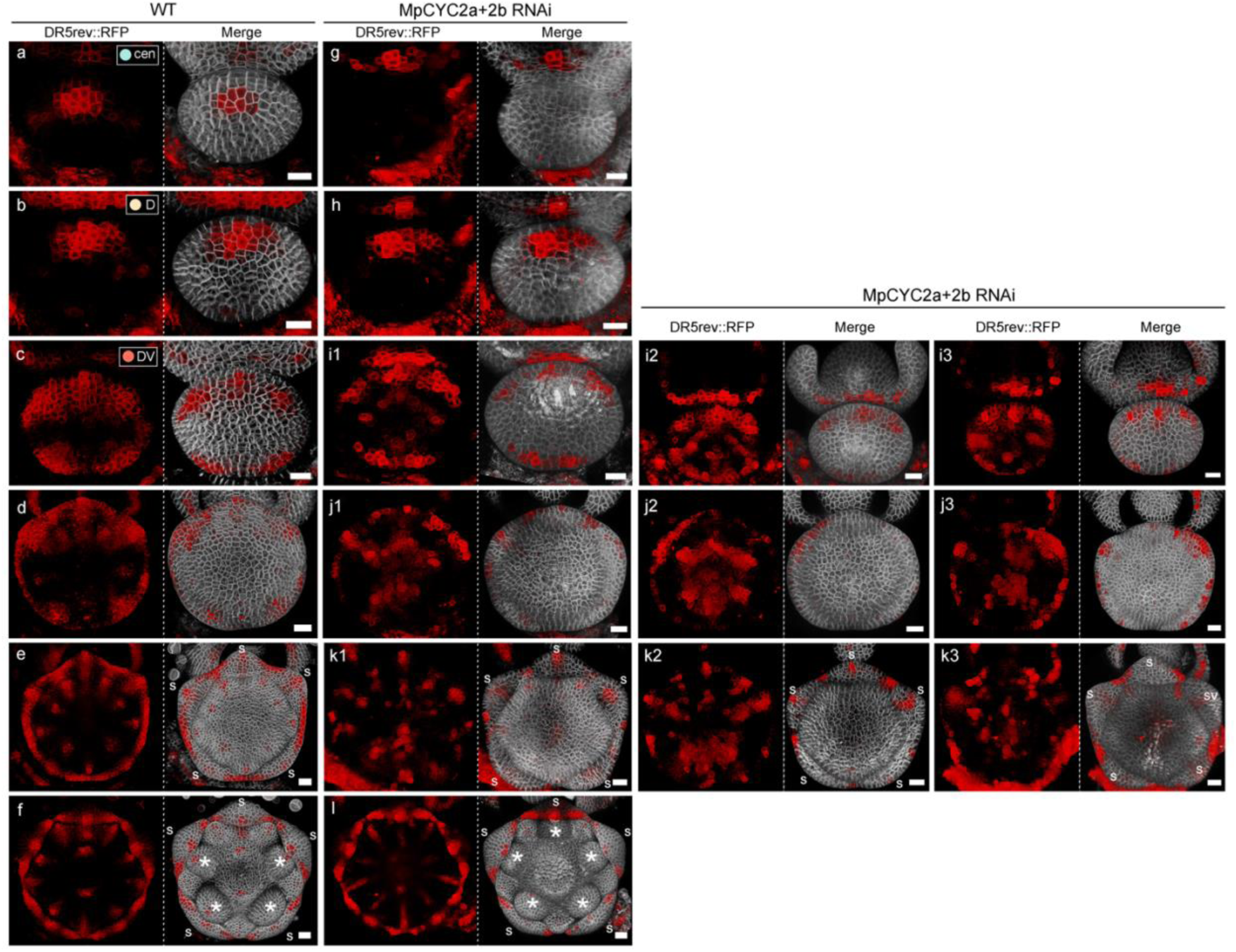
Auxin dynamics was slightly altered in MpCYC2a & 2b RNAi lines. Comparisons of DR5 patterns in WT (a-f) and *MpCYC2a*+*2b* RNAi lines (g-l) of equivalent developmental stages and sizes. (a-c, g-i): pre-organogenesis FMs. (d, j) Sepal primordia initiating. (e, k) Petal and stamen primordia initiating. (f, l) carpel primordia initiating. Panels i1-3, j1-3, and k1-3 represented three independent samples showing disorganized DR5 patterns during these stages. In comparison, the DR5 patterns in WT showing in the current figure, together with Fig. 1 and Fig.3, were very robust and reproducible at every stage. Each panel contained maximum projections of DR5rev::RFP-ER (left), merged maximum projections with plasma membrane marker (right). No DR5 was detected in the FM of RNAi lines until (h). s = sepals. Asterisks in (f, l): stamens. Scale bars = 20 µm.

**Figure S7.**
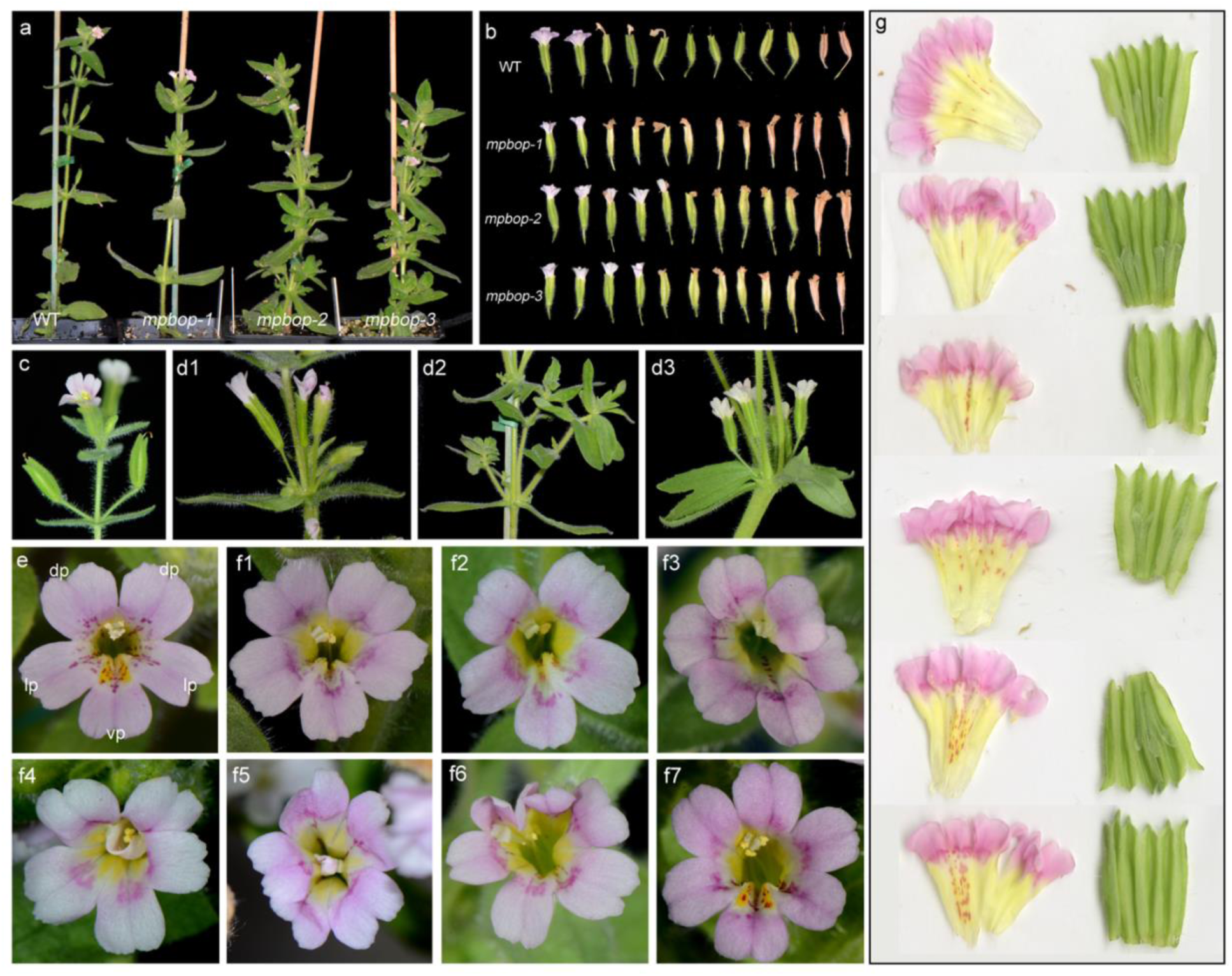
Phenotypes of mpbop plants. (a) Overall plant morphologies of a WT and three homozygous *mpbop* lines. (b) Floral organs of all *mpbop* plants failed to abscise. Each row represented five pairs of flowers of five successive nodes from young to old along a stem. Note that petals stayed on the old and dried buds in *mpbop* lines. (c) A node of a WT plant. Leaves of each node were subtending only one flower. (d1-d3) Vegetative and axillary meristem phenotypes of *mpbop* plants, including excessive activation of axillary branches (d1-d3), occasional disruption of phyllotaxy (d2), and occasionally fascinated stems (d3). (e) A WT flower. (f1-f7) Flowers of *mpbop* exhibiting a variety of phenotypes, including partial (e.g. f1-f3) or full (e.g. f4-f6) elimination of ventral petal morphological characteristics, irregularity in organ numbers (f1-f7), occasionally homeotic transformation from stamens to petals (f4, f5). (g) Unedited scans of petals and sepals of six *mpbop* flowers showing changes in petal and sepal numbers, tissue outgrowth on sepals, and all petals acquire variable degrees of dorsalization. dp = dorsal petals, lp = lateral petals; vp = ventral petal.

**Figure S8.**
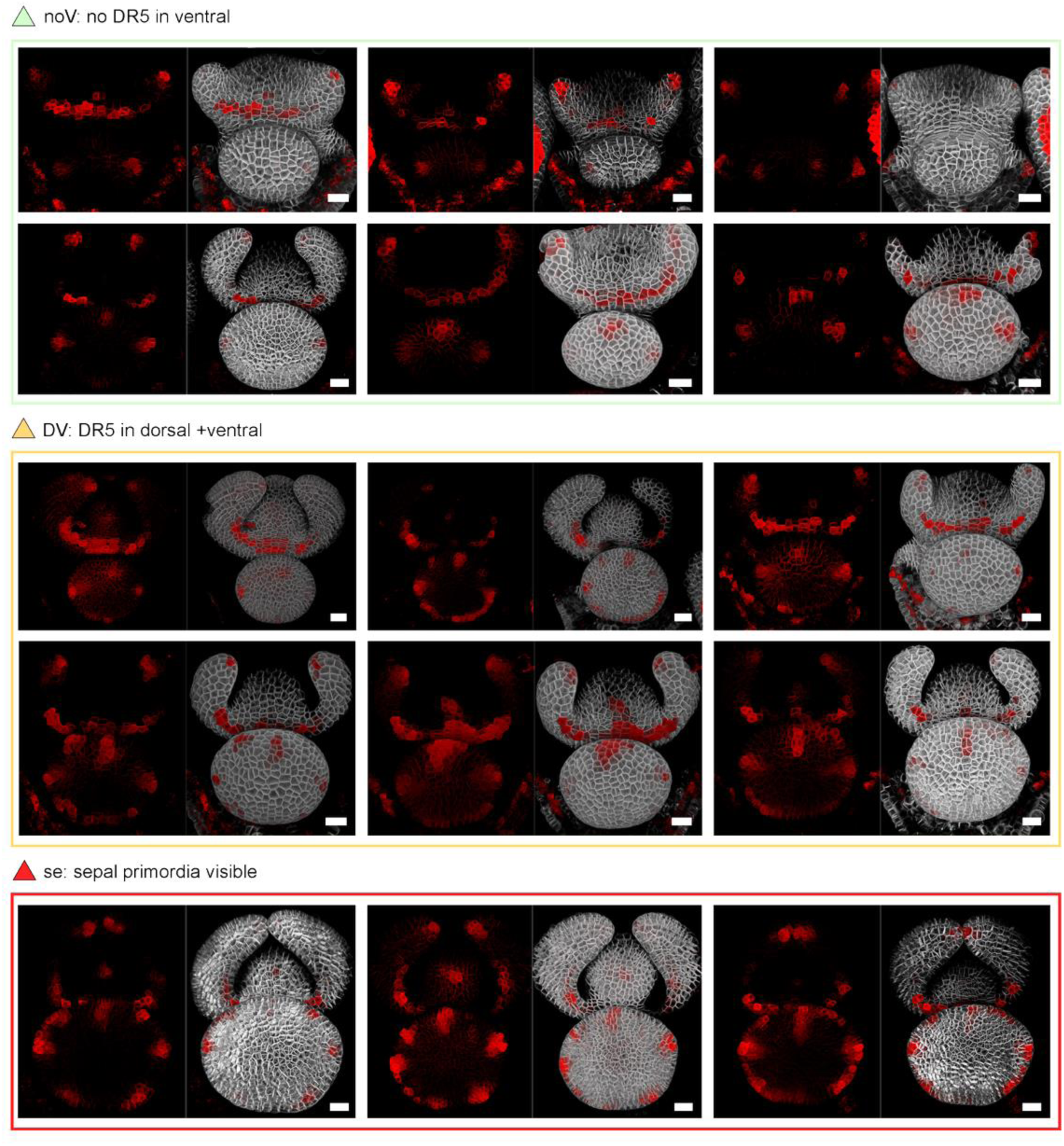
DR5 patterns in early FM morphogenesis of *mpbop*. Each panel showed maximum projections of DR5rev::RFP-ER (left) and merged with plasma membrane marker (right). For each DR5 pattern category, the locations and the number of cells expressing DR5 slightly varied from sample to sample, but the general pattern that first no DR5 in the ventral size, then DR5 in both dorsal and ventral sides, followed by sepal primordia initiate, was consistent. Scale bars = 20 µm.

**Table S1.**
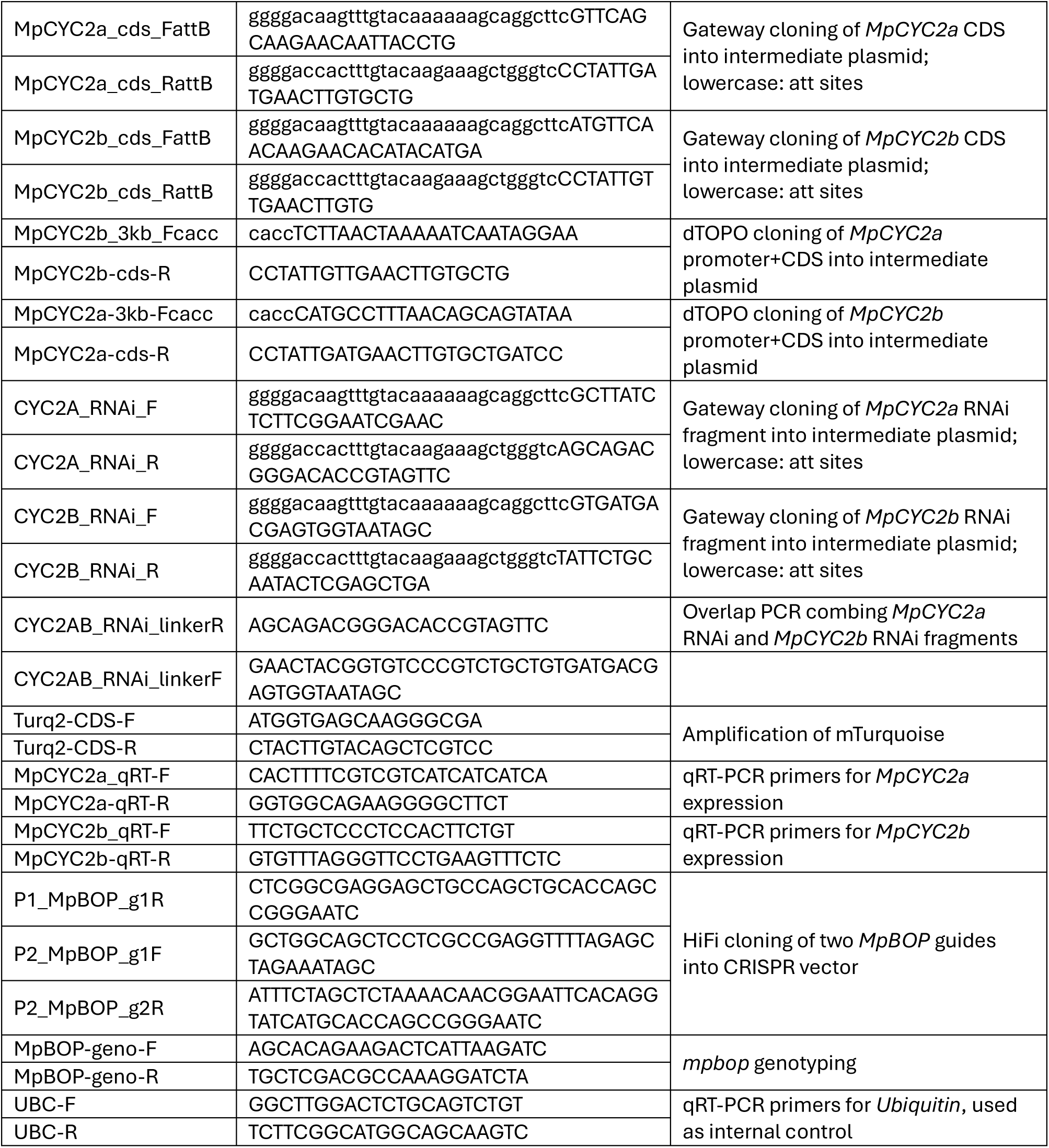
list of primers used in this study.

